# Green Fabricated Zinc Oxide Nanoformulated Media Enhanced Callus Induction and Regeneration Dynamics of *Panicum virgatum* L.

**DOI:** 10.1101/2020.03.03.974675

**Authors:** Saima Shafique, Nyla Jabeen, Khawaja Shafique Ahmad, Samra Irum, Sadaf Anwaar, Naeem Ahmad, Sadia Alam, Muhammad Ilyas, Talha Farooq Khan, Syed Zaheer Hussain

## Abstract

The current study was focused on the usage of bio synthesized zinc oxide nanoparticles to increase the tissue culture efficiency of important forage grass *Panicum virgatum*. Zinc being a micronutrient enhanced the callogenesis and regeneration efficiency of *Panicum virgatum* at different concentrations. Here, we synthesized zinc oxide nanoparticles through *Cymbopogon citratus* leaves extract to evaluate the influence of zinc oxide nanoparticles on the quality of plant regeneration in switchgrass. X-ray diffraction (XRD) and attenuated total reflectance-Fourier transform infrared (ATR-FTIR) validate phase purity of green synthesize Zinc oxide nanoparticles whereas, electron microscopy (SEM) has illustrated the average size of particle 50±4 nm with hexagonal rod like shape. Energy dispersive Xray (EDS) spectra depict major peaks of Zn (92.68%) while minor peaks refer to Oxygen (7.32%). ZnO NPs demonstrate the incredibly promising results against callogenesis. Biosynthesized ZnO NPs at optimum concentration showed very promising effect on plant regeneration ability. Both the explants, seeds and nodes used in study showed dose dependent response and upon high doses exceeding 40 mg/L the results were recorded negative, whereas at 30 mg/L both explants demonstrate 70 % and 76 % regeneration frequency. The results conclude that zinc oxide nanoparticles enhance plant growth and development. Being one of the essential plant nutrients, ZnO has greatly tailored the nutritive properties at nano-scale.

## Introduction

Switchgrass (*Panicum virgatum* L.) is a native, perennial and warm season grass, suited for biomass production for renewable fuels and also fodder on marginal soils [1]. As a perennial rhizomatous grass, it has a broad climate tolerance, rapid growth rate, high biomass yield, strong tolerance to low fertilizer level and varied abiotic and biotic stress [2]. It has strong root system and delivers unique soil preservation and is well-matched with usual farming practices [3].

Moreover, Switchgrass has less of an affinity for fertilizers [4]. As a C_4_-type, Switchgrass owns numerous agricultural benefits over C_3_-type plants and more effectual than conventional grasses, because of low nourishment requirements and less costs of harvesting [5]. Conventional breeding of switchgrass biomass is difficult because it displays self-incompatible hindrance [6]. As an undomesticated plant switchgrass has great potential for agronomic and biofuel trait improvements [7]. Switchgrass has a large genome, allopolyploidy, self-incompatibility, a long life cycle, and large stature-all suboptimal traits for rapid genetics research [8].

Using biotechnology is important to realize the potential of the biomass and biofuel-related uses of switchgrass. Tissue culture techniques aimed at rapid propagation of switchgrass have been developed [9]. With the current interest in cellulosic biofuel feedstocks, switchgrass (*Panicum virgatum* L.) research has increased in North America over the past decades [10]. So far, Tissue culture is used for the selection of different explants or plantlets and endows opportunity to study different characteristics of plant development along with growth in controlled sterilized conditions [11].

Nanotechnology have substantially changed the modern scientific era by using different methods to synthesize nano-sized particles [12]. NPs are tiny materials having size ranges from 1 to 100 nm[13 khan 2019].Nano-technology has changed the properties of metal elements’ delivery into and effect on living systems[14]. This field has vast application in different areas such as food, biomedical research, cosmetics, health care, drug delivery [15] and different domains like energy science, photochemistry have benefitted from nanotechnology [16]. Development of nanomaterials and Nano devices opened up unique application in agriculture and plant biotechnology [14]. Metallic nanoparticles have been synthesized through plethora of techniques; among that green engineering of the nanoparticles or nanomaterials is more eco-friendly and cost effective method [17]. The application of the micronutrients as fertilizer in agriculture in the form of nanoparticles has been a significant way to release necessary nutrients [18]. NPs have shown ability to boost with broad range of physiological progressions including growth and photosynthesis of plant [19].

Zinc as an essential metal for the plant growth and concentration of Zn in soil has potent effects on development and growth of plants [20]. Zinc oxide nanoparticles (ZnO NPs) enhance the development and growth of plants when incorporated or absorbed into transport system of plants and disperse in the plant cell because of the nano size [21] Translocation and absorption of NPs in various parts of plants depend upon concentration, solubility, exposure time, anatomy and bioavailability of plants [22]. Moreover, Zinc as micronutrients play key role in protein synthesis, carbohydrate metabolism, phyto-hormonal regulation particularly auxins and stress alleviated responses have been attached with sufficient delivery of zinc in plants [23].

Until now, most of the work has been done on use NPs as anti-microbial agent in tissue culture studies to provide sterilized media for explants. Keeping in view the importance of ZnO in plants growth, the current research aimed to study nutritional effect of green synthesis ZnO nanoparticles in callogenesis and morphogenesis of the *Panicum virgatum* which will provide an additional version of using NPs in the field of plant tissue culture.

## Experimental

### Extract preparation

The leaves of *Cymbopogon citratus* were collected from Kashmir and northern areas of Pakistan and identified by taxonomist experts. The fresh leaves were washed thrice with d.H_2_O to remove dust particles and dried at room temperature under shade to avoid dissociation of secondary metabolites. The dried leaves were crushed into fine powder by utilizing electrical grinder. 50g of fine leaves powder were dissolved in 500 ml of dH_2_O and boiled for 1 h at 100°C. For maximum extraction plant solution were then positioned in orbital shaking incubator for overnight at 37°C (60 rpm). Afterward extract was filtered through eight layer of muslim cloth and Whatman filter paper no.1 separately. The extract was stored at 4°C for further processing [24]

### Green Synthesis of ZnO NPs

Around 8.93 g of zinc nitrate hexahydrate salt (Sigma Aldrich) was mixed in 300 ml of the leaves extract (pH 5.7), and placed at hot plate magnetic stirrer set to 80°C for 2 hours. The reaction solution was twice centrifuge for 15 minutes at 12,000 rpm in GR-BioTek centrifuge; Greyish white pellet set down at the base of the tubes and was collected. Pellets were washed with deionized H_2_O thrice. ZnO-NPs obtained were dehydrated utilizing hot air drying oven (Memmert) at 60°C for 6 hours. Moreover, to obtain the crystalline zinc oxide nanoparticles calcinations were done at 400°C for 2 hours in Gallenkamp-furnace [25].

### Structural characterization of ZnO NPs

Structural and optical characterization of green synthesized ZnO NPs were performed to evaluate their diameter, purity, surface modification and configuration through Scanning Electron Microscope (SEM,TESCAN, MIRA3), Energy Dispersive Spectroscopy (EDS,Oxford), X-ray Diffraction (XRD, GNR Analytical Instruments Groups) and ATR-Fourier Transform Infrared Spectroscopy (ATR-FTIR, Thermoscientific) analysis. Structural along with elemental aspects of the green-synthesised ZnO-NPs was characterized using scanning SEM examination with JOEL JSM 6490LASEM operating at accelerating voltage of 20 kV along EDX detector. Crystalline structure of powder NPS was assessed through XRD. Crystallographic structure of green-synthesized ZnO NPs were obtained utilizing copper (Cu)-Kα radiation [λ = 1.54060 Å] with nickel monochromator in the range of 2θ between 20° and 80°. Scherrer’s formula is used to standardize average crystallite size. Furthermore, to identify vibrational characterization of green engineered ZnO NPs, and functional groups engage in reduction or capping of ZnO NPs were assessed using ATR-FTIR spectroscopy within the wave number varying from 500–4000 nm.

### Collection of explant and optimization of culture conditions

Mature plants and seeds of Switchgrass were collected from NARC (National Agriculture Research Centre). Seeds from mature spikelet’s were dehusked and taken as explants along with internodes of fresh stems. The explants were surface sterilized following protocol of [26]. MS (Murashige and Skoog) 4.3 g/L media with 3% sucrose, 10 ml (100 XL) of Vitamin B12, 3.5 g/L of casein hydro lysate and 4g/L gelrite agar was mixed, Afterward pH of media was maintained at 5.8 and autoclaved the media for 20 min at 121°C. Different concentration ZnO-NPs were added in autoclaved media. For prevention of agglutination of nanoparticles in base of media, the media was set to normal up to 45°C and media tubes were kept at room temperature to solidify. All the work was carried out under sterilized conditions.

### Callus Induction

For the induction of callogenesis, MS media with 2.5 mg/L of 2,4-D was utilized and augmented along with different concentration of ZnO-NPs (10–50mg/L), and without ZnO-NPs as control[27]. The culture tubes were placed in dark as well as in light (16/8h photoperiod) conditions at 23 ± 2°C for 2 weeks. After regeneration period of 2 weeks, embryogenic callus was attained. The frequency of callus induction was recorded by using following formula:

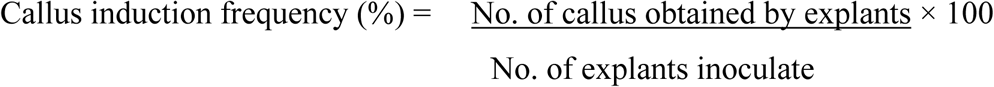

### Callus Regeneration

Embroyogenic callus formed were shifted on the regeneration media RGM (2.5 mg/L of BAP + 0.5 mg/L of Kin) with varying concentrations of ZnO-NPs and without ZnO-NPs as control. Different concentrations of ZnO NPs were supplemented to obtained best regeneration frequency of switch grass. After 10–12 weeks regeneration frequency was calculated by following formula:

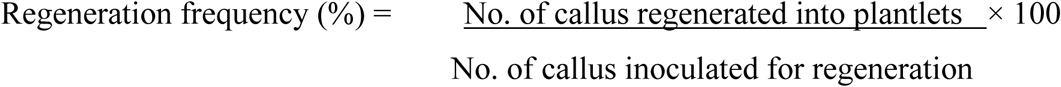

### Statistical analysis

Data was statistically analyzed by two factorial design and Analysis of variance (ANOVA) in Statistix 8.1 software. All experiments were replicated thrice and individual replication had 10 treatments.

## Results and discussion

### SEM Analysis

The morphology of biologically synthesized ZnO was studied using Field Emission Scanning Electron Microscope (FE-SEM) while its chemical composition was determined using energy dispersive x-ray spectroscopy (EDS), as depicted in Fig.3. ZnO prepared through green synthesis had a hexagonal shape and a rod-like morphology, so these will be referred to as ZnO nanorods. Fig.2. shows the SEM images of ZnO nanorods at increasing magnification of 5μm, 2μm, 1μm and 500nm, respectively, and individual ZnO nanorods can be clearly noticed at higher magnifications. The diameter of ZnO nanorods was found to vary in the range between 90-390nm. EDS in Fig. 2(e) shows that the ZnO nanorods are of high purity since only two peaks were observed in the EDS spectrum. The major peak for Zn indicates 92.68 wt% Zn while the minor one refers to the remaining oxygen content (Table 1). EDS results showed that Zn and O ions are present in the green synthesized NPs. The current observations are supported by findings of [28]. [29] reported similar hexagonal shape of the NPs. Modifying the pH results in alteration of surface charge of phytochemicals which affects their reduction capacity and binding potential of metal ions in the synthesis of NPs [30]. In our results Hexagonal nano rods like morphology of synthesized NPs utilizing SEM clearly indicates victorious capping of bio active compounds from *C. citratus* extracts.

**Table 1.** Atomic percentage of Oxygen and Oxygen in Zinc oxide NPs as observed from EDX spectra.

| Element | Weight% | Atomic% |
| --- | --- | --- |
| Hexagonal ZnO NPs |  |  |
| Peaks possibly omitted : | 0.270, 2.149 keV |  |
| O K | 7.32 | 24.40 |
| Zn K | 92.68 | 75.60 |
| Totals | 100.00 |  |

**Fig.1.**
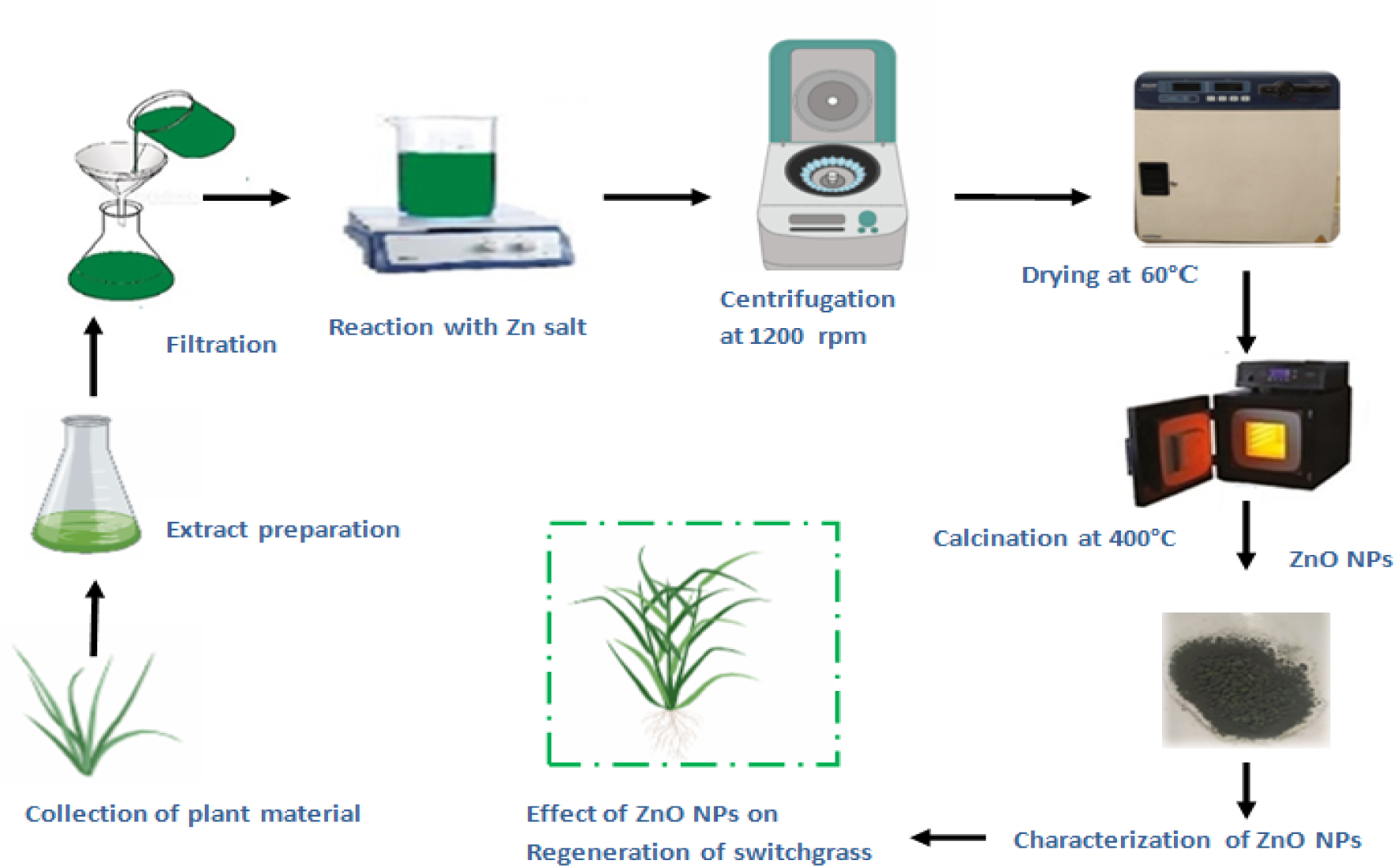
Schematic diagram of green-synthesized of ZnO-NPs using *Cymbopogon Citratus*.

**Fig. 2.**
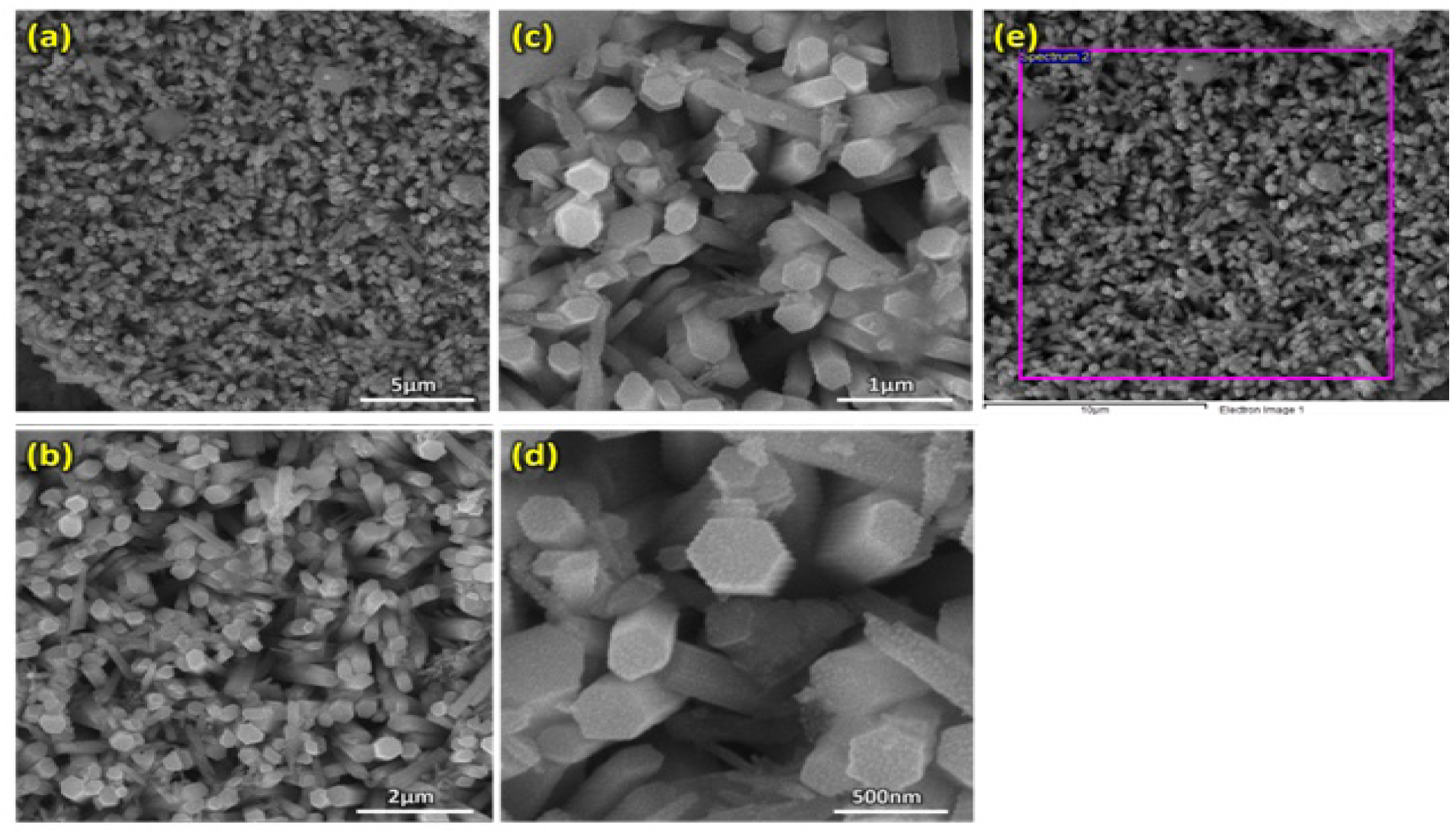
FE-SEM images of ZnO nanorods at a resolution of (a) 10KX, (b) 25KX, (c) 50KX, (d) 100KX, and (e) EDS spectra of ZnO nanorods.

**Fig. 3.**
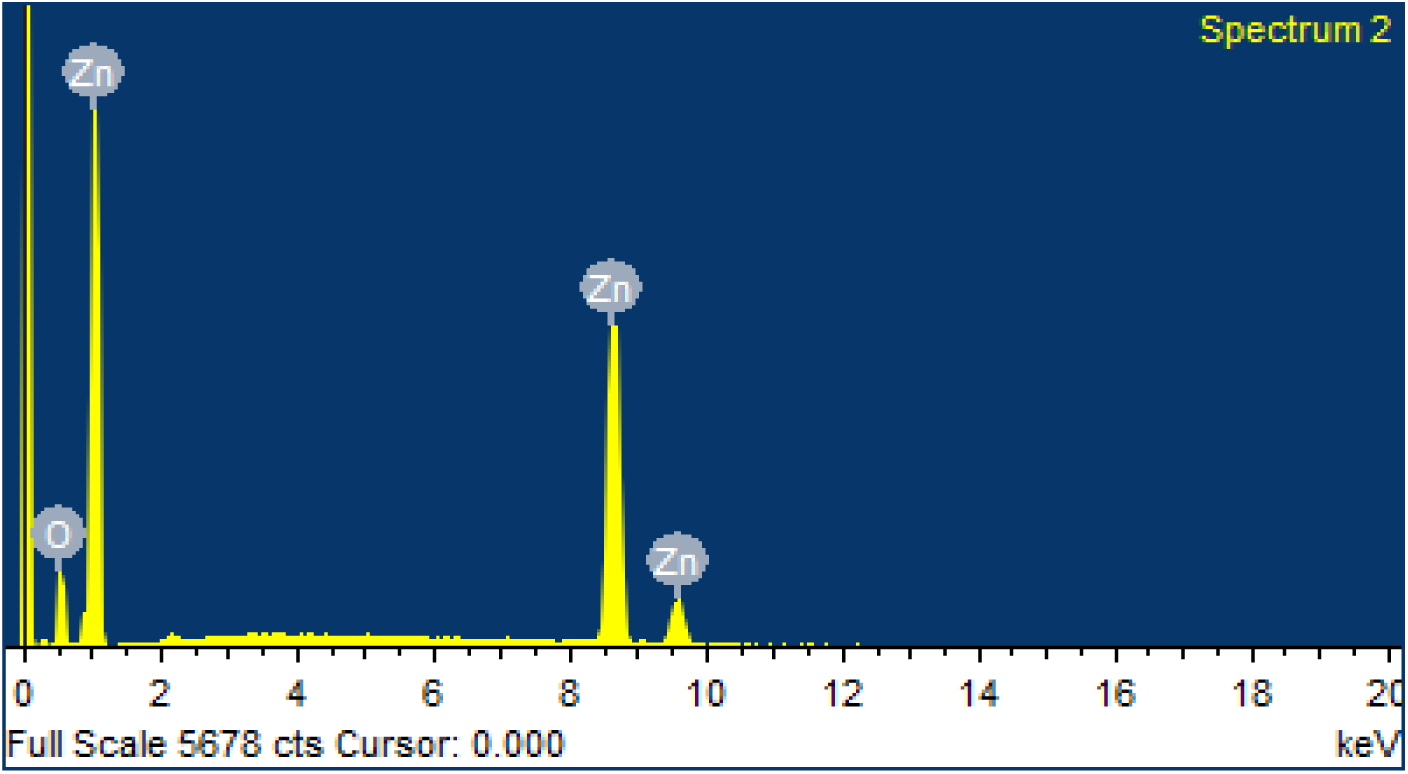
EDS analysis of ZnO NPs indicating the purity of Zinc (Zn) and Oxygen (O) in the sample.

### FTIR Spectroscopy

The functional groups present in ZnO NPs and plant extracts were interrogated through ATR-FTIR in the range of wavelength number 500-4000cm^-1^ (Fig.4). ATR-FTIR spectra of ZnO results in peaks range between 3852.85, 3649.37, 2167.06, 2036.06, 2011.69, 1980.83, 1043.68. 794.82, 746.04, 727.34, 677.37, 668.90, 660.07 and 652.81 cm^-1^.The sharp peak at 2167.06 cm^-1^ indicate the stretching vibration of C≡C stretch of Alkynes.The sharp peak at 1043.68 cm^-1^ shows the presence of amine NH group and two peaks at 1043.68 cm^-1^ refers to the presence of C-H stretch of aliphatic amines [31]. The peaks in the region below 700cm^-1^ are assigned to Zn-O which shows ZnO NPs absorption band near 660cm^-1^. Vibrational and stretching nature of molecules in FTIR spectra provide data to examine the pure phase of ZnO NPs. Zinc oxide demonstrate typical identified FTIR spectrum (Fig.5) and results demonstrate the secondary metabolites attachment with ZnO NPs. Electrostatic forces between the positive charged Zn ions and negatively charged molecules are involve in binding of these functional groups. This interface makes NPs ideal applicant for different biological activities [32].

**Fig. 4.**
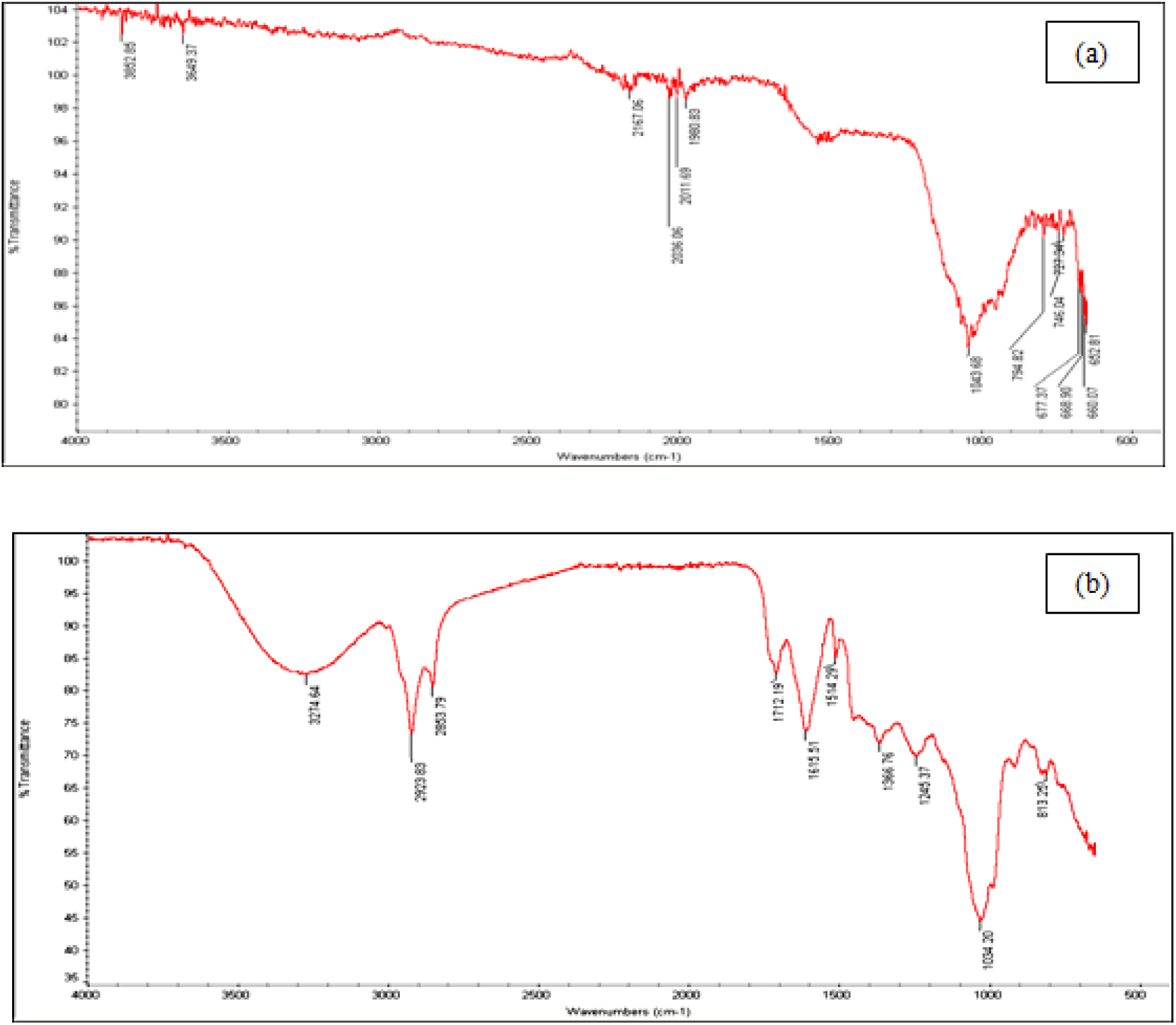
ATR-FTIR spectrum of the powder samples (a) green synthesized ZnO NPs (b) *Cymbopogon citratus* extracts.

### XRD Analysis

The phase purity and crystalline nature of NPs has been investigated through XRD. Fig. 5. illustrate crystallographic nature of green synthesized NPs in the range of 20-80 The XRD pattern shows clear and distinguishable peaks at (100), (002), (101), 102, (110), (103), (200), (112), (201), (004) and (202) corresponds to 2θ values of 31.73, 34.43, 36.21, 47.55, 56.56, 62.82, 63.45, 65.21, and 69.02, 70.02, 75.45, 77.05. The comparatively high amount of the (101) peak is revealing of anisotropic development and suggests a ideal orientation of the crystallites. All noted peaks intensity profiles were features of the hexagonal rod structure. The structural crystalline size of ZnO NPs is calculated using Scherrer’s formula;

**Fig. 5.**
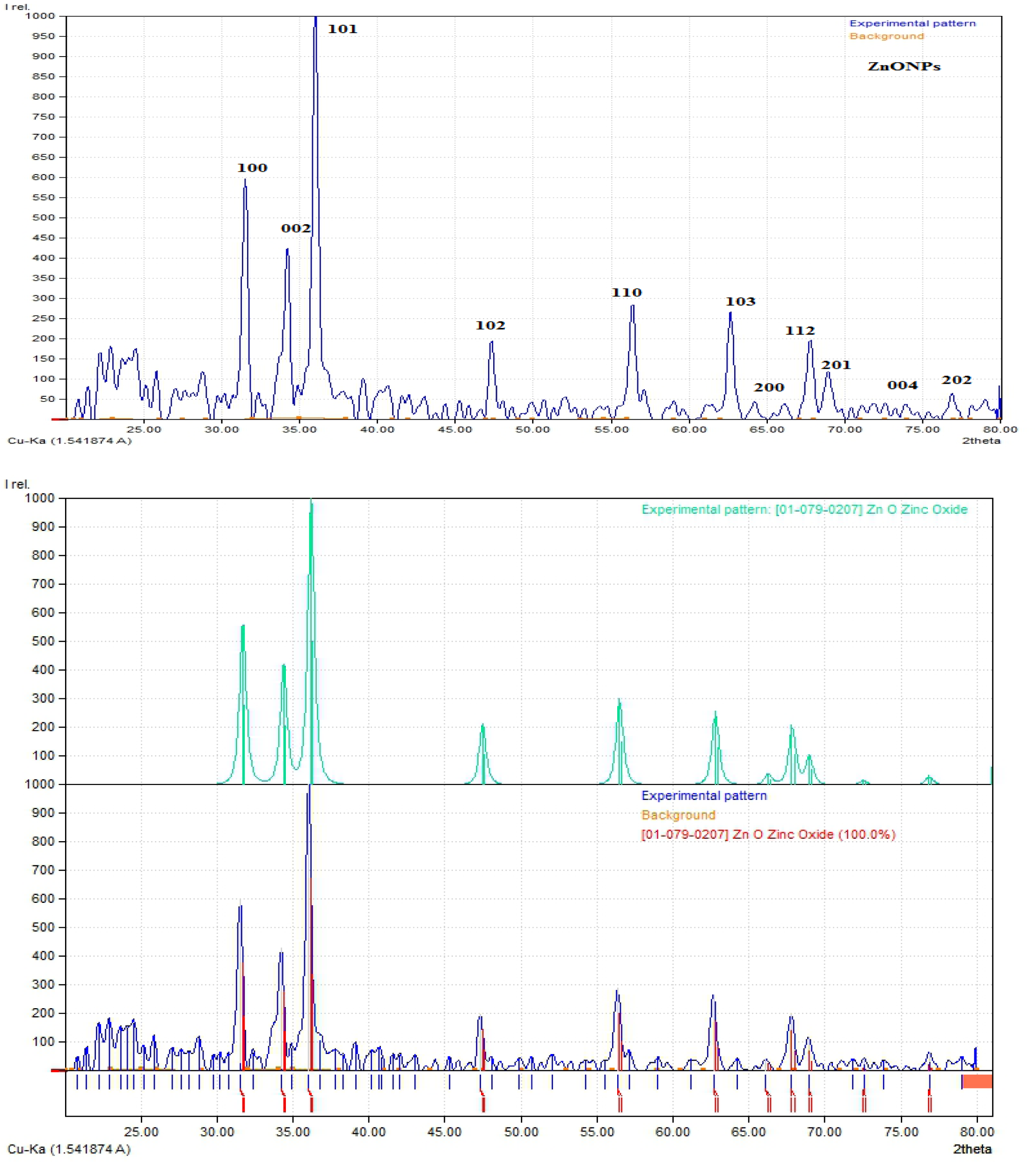
XRD pattern of green synthesized Zinc oxide nanoparticles.

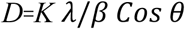

K is shape factor, λ is the wavelength of X-ray, and β and θ are the half width of the peak and half of the Bragg angle correspondingly. The average crystalline size of are found closer to 50nm. All the peaks in XRD specta are accordance to JCPDS card no. 00-004-0347 which validate the hexagonal crystalline structure and no peaks relate to any type of impurity.

### 3.4. Effect of ZnO-NPs on MS Medium for Callus Induction Frequency

In present study the role of green synthesized ZnO NPs from C. Citratus was investigated for its efficacy in tissue culture of Switchgrass. This is the first reported study on switchgrass tissue culture using ZnO NPs. Zn also play a vital role as it is a co factor for appropriate functionality of several enzymes comprising superoxide catalase and dismutase, which inhibits ROS stress in plant cells [33]. The role of ZnO-NPs in induction of callus was investigated in the present study and data is presented in Table 2. The tested concentrations of NPs were 10, 20, 30, 40 and 50 mg/L (Fig. 6). At concentration 10 mg/L there was an increase of callus induction frequency 63% as compared to 53% in control. At concentration 30 mg/L, seeds showed maximum callogenesis of 90% followed by 83% at concentration 20mg/L as compared to control. Internodes as explants showed similar results against tested concentrations but the maximum callogenesis 96% was recorded at 20 mg/L and at 30 mg/L it was 86% as compared to control 63%. Metal oxide NPs like Silver, Zinc oxide, copper oxide and cerium oxide have influences on the plants growth [34]. When the concentration of nanoparticles was further increased to 40 mg/L there was a decreased in callus induction frequency in both seeds and internodes to 43% and 5% respectively. The results showed very low callogenesis 16% at 50 mg/L in internodes but no callus was recorded at that concentration in seeds as explants.. The increase in callus induction frequency using ZnO NPs can be explained in terms as Zn is a micronutrient and can be supplied to plants NPs [25], However the effects were concentration dependant and above 20-30 mg/L was proved to increase the callogenesis. Data clearly reflects the positive response of using ZnO-NPs at low concentration upto 30 mg/L. As the concentration increased from 30mg/L the effects of ZnO NPs were shifted towards negative response due to the injury in the cell’s wall and membrane thus piercing it and interfering with the various plant’s developments [35]. As they can restrictthe electron transport chain of mitochondria and chloroplast, which may cause oxidative burst, and rise in ROS concentration and cause cell death [36] which in turn decrease the collogenesis in swtichgrass.

**Table 2:**
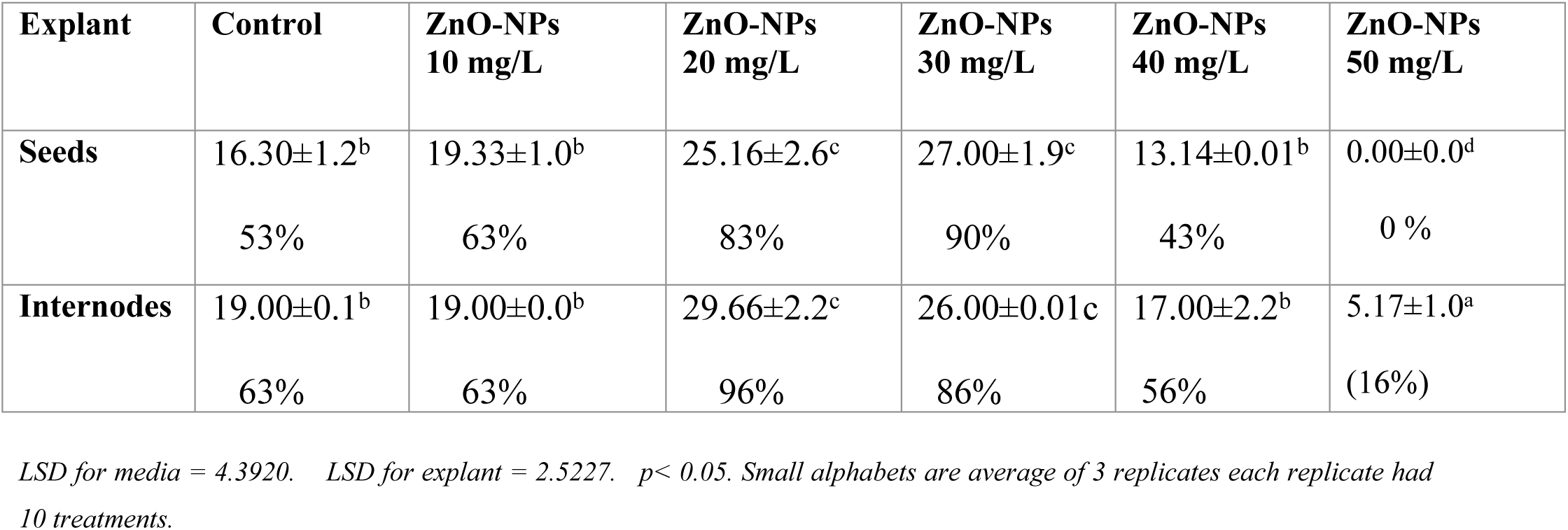
Effect of Various Concentrations of ZnO-NPs on MS Medium for Callus Induction Frequency:

**Fig.6.**
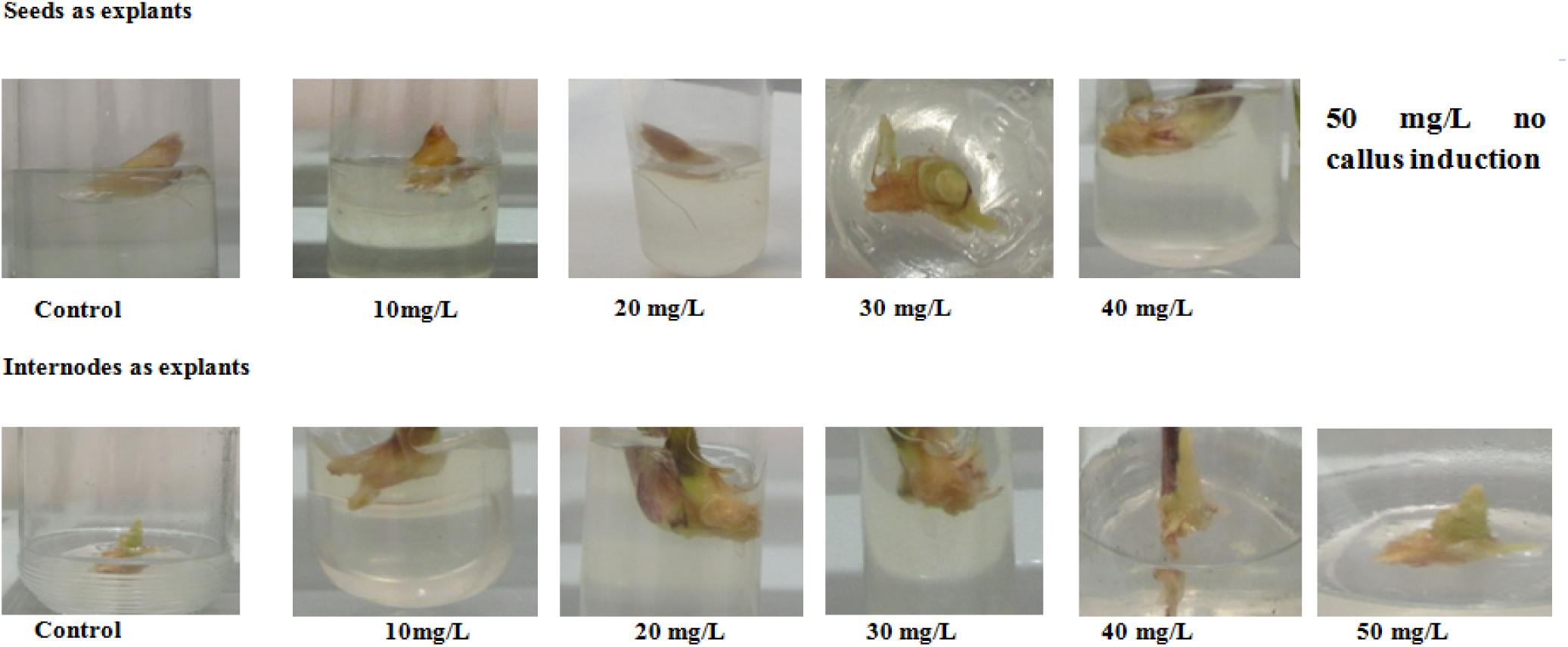
Effect of different concentration of ZnO NPs (10, 20, 30, 40, 50 mg/L) on callogenesis.

In tissue culture media the enhanced growth frequency of callogenesisis may be due to up regulation or down regulation of different hormonal pathways. The increased level of cytokinin in response to NPs and enhanced rate of callogenesis [37]. There is a difference in response of both explants tested in terms of optimum growth at different concentrations of NPs. Hence, it can be concluded that nanoparticles having small size can enter the explants, and thus affect some genetic reprogramming features [38].

### 3.5. Average Number of Days to Callus Initiation

Number of days explants took to start callogenesis was observed and recorded in Table 3. Both the explants showed variability in terms of number of days for callus initiation at various concentrations of ZnO-NPs as compared to control 29 days. At concentration 20 mg/L seeds took minimum of 19 days, followed by 30 mg/L at which callogenesis was observed after 20 days of inoculation. However, with the increase of concentration there was an increase in days for callus initiation at concentration 40mg/L as compared to control. At 10 mg/L no significant difference was observed in both the explants in starting callus as compared to control and no callogenesis was recorded when concentration increased to 50mg/L.

**Table 3:** Average Number of Days to Callus Initiation

| Explant | Control | ZnO-NPs<br>10 mg/L | ZnO-NPs<br>20 mg/L | ZnO-NPs<br>30 mg/L | ZnO-NPs<br>40 mg/L | ZnO-NPs<br>(50 mg/L) |
| --- | --- | --- | --- | --- | --- | --- |
| **Seeds** | 29±1.45 | 27±3.22 | 20±2.15 | 20±0.01 | 32±0.76 | _____ |
| **Internodes** | 23±1.05 | 23±0.45 | 17±2.75 | 15±1.85 | 29±0.98 | 33±1.55 |

In case of internodes the best results were observed at concentration 20mg/L and callogenesis started at 15 days as compared to control 23 days followed by 17 days at concentration 20mg/L. No difference in initiation of callus was observed at minimum tested concentration of NPs i-e 10 mg/L as compared to control both took 23 days. Maximum 29 and 33 days was recorded for the concentrations 40mg/L and 50mg.L respectively. As the callus induction frequency was declined with increase in concentration and more days were taken to initiate callus formations. This is because of the negative effects or cytotoxicity and genotoxicity of ZnO at higher concentration [39].

Zinc shows significant role in a broad range of processes in plants, like growth hormone synthesis and internodes elongation. The optimum concentration at which minimum days taken by Internodes were17 days as compared to control explants which took 23 days to initiate callus. This phenomenon can be explained in terms of up regulation of genes, as the nano sized ZnO NPs can easily penetrates into cytosol via active transport and can also help in regulation of other processes like cell signalling, recovering and the regulation of plasma membrane [40]. Plants grow unorganized cell masses such as callus and tumours in reaction to various stress inducements like NPs as overexpression of genes encoding auxins cause proliferation of cells that leads to initiate callogenesis [41].

### Regeneration of Switchgrass at different concentrations of ZnO-NPs

The calli resulting from both the tested explants were shifted to regeneration media control and with various concentrations of ZnO-NPs to check the effectiveness of NPs on regeneration frequency reported on RGM with 20mg/L and 30mg/L ZnO-NPs in internodes and seeds respectively. After shifting to RGM media callus were regenerated within few days (Fig.7) as these NPs interact with plants and became a source of change in physiology and morphology of plants [42]. The regeneration capacity of switch grass was enhanced by addition of ZnO NPs as compare to control (Table 4).The effects of NPs mainly due to arrangement, size, and concentration [43]. Small size to higher surface area to charge ratio enables these particles to dissociate quickly in the cytosol of the plant and because of this speedy release of ZnO-NPs aids enzymes at cellullar level to achieve precise and effectual photosynthesis process [44]. Our findings are in accordance with increase the concentration of ZnO NPs enhance regeneration in Banana plant reported by [45]. [46] reported the effect of ZnO on *Arachishypogaea* growth up to 1,000 mg/L the NP promoted seed germination and growth dynamism.

**Fig. 7.**
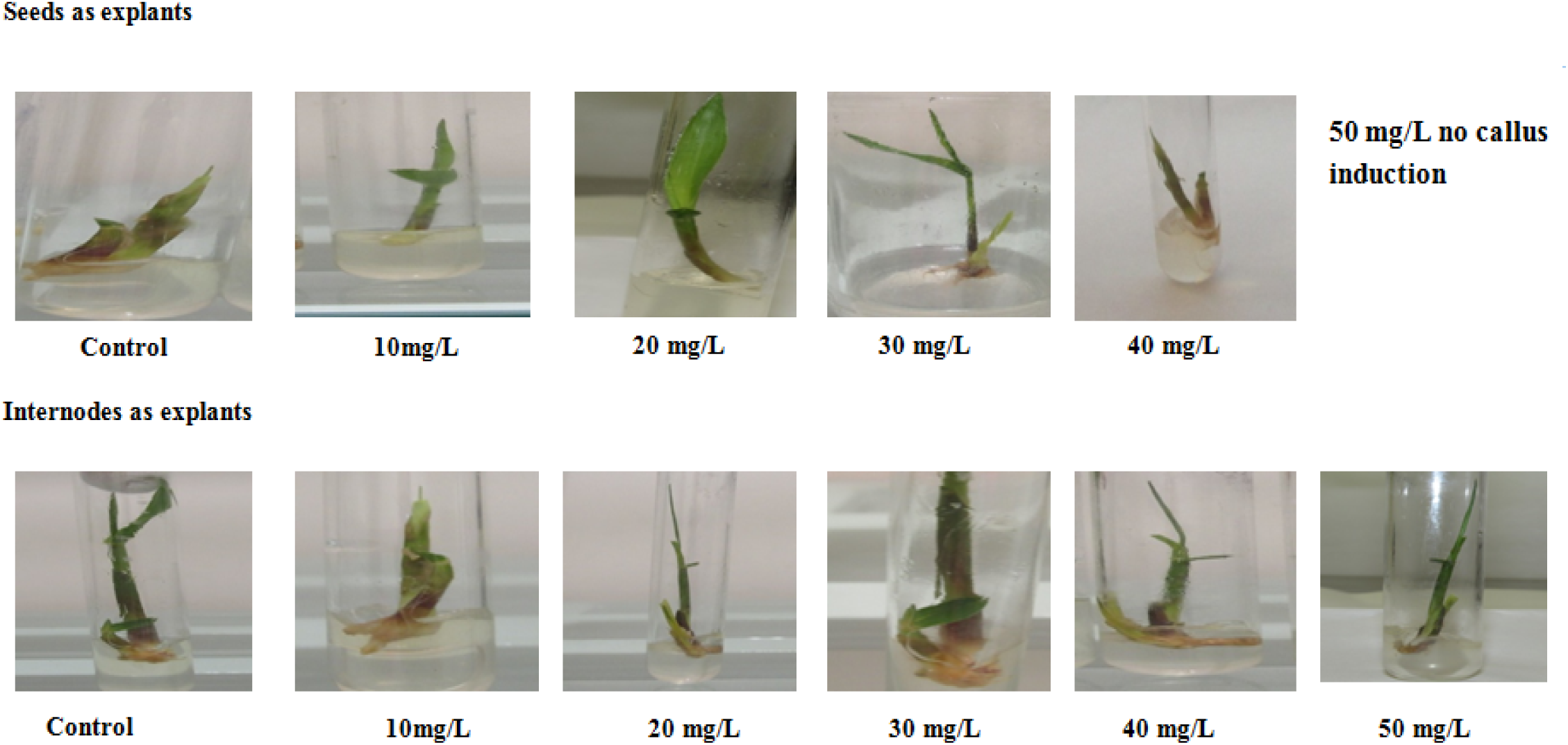
Regeneration frequency of Switchgrass at different concentrations of ZnO NPs (10-50 mg/L).

**Table 4:** Regeneration of Switchgrass at different concentrations of ZnO-NPs.

| Explant | Control RGM | RGM +ZnO-NPs<br>10 mg/L | RGM+ZnO-NPs<br>20 mg/L | RGM+ZnO-NPs<br>30 mg/L | RGM+ZnO-NPs<br>40 mg/L | RGM+ZnO-NPs<br>50 mg/L |
| --- | --- | --- | --- | --- | --- | --- |
| Seeds | 7.13±1.05<sup>c</sup> | 13.20±1.14<sup>b</sup> | 17.28±0.75<sup>b</sup> | 23.10±2.1<sup>a</sup> | 5.21±2.45 | 0.00±0.00<sup>d</sup> |
|  | 24% | 43% | 57% | 76% | 16% | 0 % |
| Internodes | 10.00±1.35<sup>c</sup> | 15.00±0.05<sup>b</sup> | 24.00±0.01<sup>a</sup> | 21.33±1.1<sup>a</sup> | 17.00±1.25<sup>b</sup> | 7.43±0.98<sup>c</sup> |
|  | 33% | 50% | 80% | 70% | 56% | (23%) |

LSD for explant= 4.019. $p < 0.05$ . Small alphabets are average of 3 replicates each replicate had 10 treatments.

The regeneration frequency can be increase by adding ZnO-NPs, as zinc helps the plant to make chlorophyll. During the process of photosynthesis most of the enzymes needed metallic ions as co-factor to complete their appropriate function whereas; accessibility and reactivity of these ions at the level of cell are vital signs for usual plant development [47]. Additionally, Zn deficiency hindered the plant bicarbonate use capability and PS II activity due to the incidence of excess bicarbonate ions. Zn deficiency represses other Zn-containing enzymes, such as alcohol dehydrogenase and glutamate dehydrogenase, consequently inhibiting plant growth [48].

However, at higher concentrations of metallic NPs growth factors showed a noteworthy decrease [49]. NPs can have considerable negative effects, like decrease in seed germination and decrease in plant growth and development resulting in wilting of plant [50]. Due to the interaction of NPs at cellular level ROS formed which interacts with nearly all cellular mechanisms resulting in protein modifications, lipid peroxidation, and destruction to DNA which finally results in necrosis or cell death [37]. Hence, optimization of concentration of NPs is required to obtain beneficial results in plant tissue culture.

## 2. Conclusion

Nanoparticles are widely used in human life as they have contributed in almost every domain of life. But there is a need of investigation to recognise the molecular mechanism of plant nanoparticle interaction so that the metallic NPs like ZnO can be used efficiently in plant tissue culture and agricultural practices. The small size and biocompatibility with plant cell cultures make nanoparticles an ideal candidate to use as bio fertilizer. Zn can be used to enhance callogenesis and regeneration of switchgrass as tissue culture is the backbone of plant biotechnology and plant genetic engineering. In future these findings will prove very helpful in modifying switchgrass with a foreign gene and regenerating through tissue culture.

